# Identification of Fluoxetine as a direct NLRP3 inhibitor to treat atrophic macular degeneration

**DOI:** 10.1101/2021.01.11.425135

**Authors:** Meenakshi Ambati, Ivana Apicella, Siddharth Narendran, Shao-bin Wang, Hannah Leung, Felipe Pereira, Akhil Varshney, Kirstie L. Baker, Kenneth M. Marion, Mehrdad Shadmehr, Cliff I. Stains, Srinivas R. Sadda, Ethan W. Taylor, S. Scott Sutton, Brian C. Werner, Joseph Magagnoli, Bradley D. Gelfand

**Author notes:** M.A., I.A., and S.N. contributed equally to this work. To whom correspondence may be addressed: Bradley D. Gelfand, Joseph Magagnoli, Brian C. Werner.

## Abstract

The atrophic form of age-related macular degeneration (dry AMD) affects nearly 200 million people worldwide. There is no FDA-approved therapy for this disease, which is the leading cause of irreversible blindness among people over 50 years of age. Vision loss in dry AMD results from degeneration of the retinal pigmented epithelium (RPE). RPE cell death is driven in part by accumulation of *Alu* RNAs, which are noncoding transcripts of a human retrotransposon. *Alu* RNA induces RPE degeneration by activating the NLRP3-ASC inflammasome. We report that fluoxetine, an FDA-approved drug for treating clinical depression, binds NLRP3 in silico, in vitro, and in vivo, and that it inhibits activation of the NLRP3-ASC inflammasome in RPE cells and macrophages, two critical cell types in dry AMD. We also demonstrate that fluoxetine, unlike several other anti-depressant drugs, reduces *Alu* RNA-induced RPE degeneration in mice. Finally, by analyzing two health insurance databases comprising more than 100 million Americans, we report a reduced hazard of developing dry AMD among patients with depression who were treated with fluoxetine. Collectively, these studies triangulate to link fluoxetine as a potential drug repurposing candidate for a major unmet medical need that causes blindness in millions of people in the United States and across the world.

**Significance Statement:** Dry age-related macular degeneration (AMD) affects the vision of millions of people worldwide. There is currently no FDA-approved treatment for dry AMD. The inflammasome components NLRP3 and ASC have been implicated in the pathogenesis of dry AMD. We report that fluoxetine, which is approved for the treatment of clinical depression, directly binds the NLRP3 protein and prevents the assembly and activation of the NLRP3-ASC inflammasome. As a result, it also blocks the degeneration of retinal pigmented epithelium (RPE) cells in an animal model of dry AMD. Furthermore, we demonstrate through an analysis of health insurance databases that use of this FDA-approved anti-depressant drug is associated with reduced incidence of dry AMD. These studies identify that fluoxetine is a potential repurposing candidate for AMD, a prevalent cause of blindness.

## INTRODUCTION

Age-related macular degeneration (AMD) is the leading cause of irreversible blindness among those over 50 years of age around the world (Ambati et al. 2003). The dry form of AMD is characterized by degeneration of the retinal pigmented epithelium (RPE), a specialized monolayer of cells lying external to the retinal photoreceptors (Ambati & Fowler, 2012). Progressive RPE degeneration in the central portion of the retina known as the macula leads to photoreceptor cell death and consequent vision loss over several years (Shen et al., 2020). Dry AMD, which accounts for approximately 90% of the 200 million global cases of AMD (Wong et al., 2014), has no FDA-approved therapy (Mitchell et al., 2018).

In dry AMD, areas of RPE degeneration display abnormal accumulation of *Alu* RNAs (Kaneko et al., 2011), which are noncoding RNAs transcribed from the highly abundant family of *Alu* repetitive elements in the human genome (Kazazian & Moran, 2017). These *Alu* RNAs as well as the related mouse retrotransposon B2 RNAs are cytotoxic (Kaneko et al., 2011; Dridi et al., 2012), as they activate the NLRP3-ASC inflammasome (Tarallo et al., 2012), a multiprotein complex that acts as a cellular danger sensor that responds to a diverse set of inflammatory stimuli (Latz, 2010; Ambati et al. 2013). In response to various danger signals, the proteins NLRP3 (nucleotide binding domain, leucine-rich repeat receptor, and pyrin domain-containing protein 3), ASC (apoptosis-associated speck-like protein containing a caspase recruitment domain), and pro-caspase-1 assemble into a macromolecular platform known as the ASC speck (Masumoto et al., 1999). The defining molecular event of inflammasome activation is autocleavage of pro-caspase-1 into active caspase-1. Active caspase-1, in turn, enzymatically cleaves two interleukins (ILs), IL-1β and IL-18, from their inactive pro-form to their mature, active forms. Active IL-1β and IL-18, which are elevated in dry AMD macrophages and RPE cells, respectively, are pro-inflammatory cytokines. In dry AMD, activation of this inflammasome occurs both in RPE cells (Tarallo et al., 2012) and in macrophages (Eandi et al., 2016), and leads to retinal cell death.

Despite dozens of clinical trials over two decades, no treatment has yet proven effective for dry AMD (Mitchell et al., 2018). We sought to identify a novel therapy for dry AMD by employing the concept of drug repurposing (Boguski et al., 2009). Specifically, we hypothesized that an existing drug that is FDA-approved for another disease and that shares structural similarity to a known inflammasome inhibitor might be effective against dry AMD.

Here we demonstrate that fluoxetine, which is FDA-approved for major depressive disorder, contains a structural moiety present in a known NLRP3 inhibitor and that it directly interacts with NLRP3 and inhibits its assembly and activation. Further, fluoxetine inhibits *Alu* RNA-induced RPE degeneration in mice. We also present evidence from two health insurance databases that fluoxetine use is associated with reduced incident dry AMD, suggesting that it potentially could be repurposed.

## RESULTS

### Fluoxetine interacts with NLRP3

The small molecule CY-09 is reported to inhibit the NLRP3 inflammasome (Jiang et al., 2017). By examining the structure of various FDA-approved drugs, we observed that fluoxetine, which is indicated for clinical depression, shared structural similarity to CY-09 in that both possessed a (trifluoromethyl)phenyl moiety (Fig. 1*A*).

**Figure 1.**
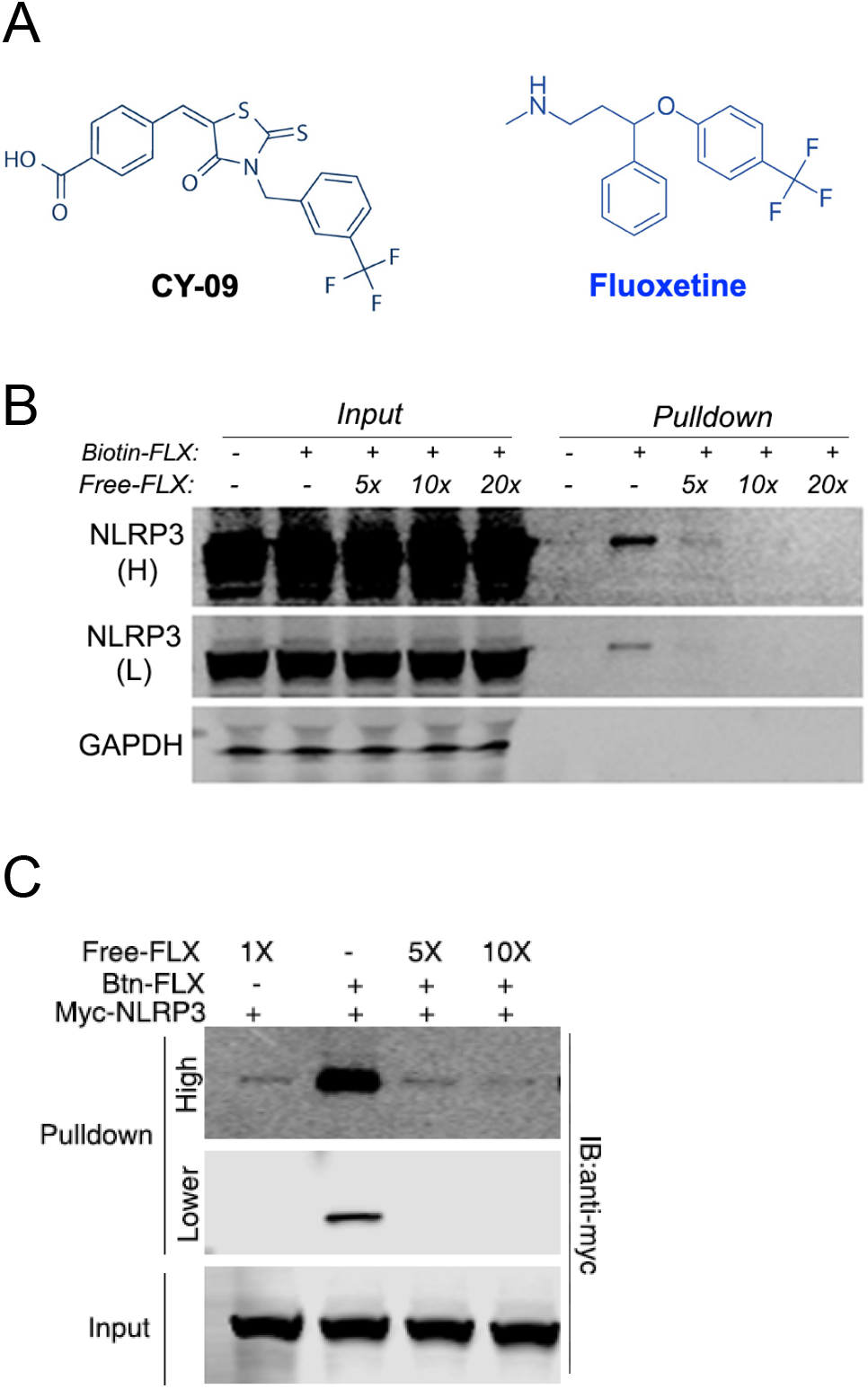
Fluoxetine binds NLRP3. **A.** CY-09, a small molecule inhibitor of NLRP3 (Jiang et al., 2017), and fluoxetine both contain a (trifluoromethyl)phenyl moiety. **B.** Biotinylated fluoxetine was incubated with protein lysate collected from LPS-primed human THP-1 monocytes. Protein complexes were precipitated by streptavidin pulldown. The input and pull-down fractions were immunoblotted for NLRP3 or GAPDH. High (H) and low (L) exposures shown. **C.** In vitro interaction of biotinylated fluoxetine (Btn-FLX) and purified Myc-NLRP3 protein analyzed by streptavidin pulldown. Excess amounts (5–10×) of free (non-biotinylated) fluoxetine were used as competitors. High and lower exposures shown.

To test whether fluoxetine binds NLRP3, we synthesized a biotin conjugate of fluoxetine (*SI Appendix*, Fig. S1 *A–C*). We then incubated biotinylated fluoxetine with protein lysate collected from LPS-primed human THP-1 monocytes. Fluoxetine-associated protein complexes were precipitated by streptavidin pull-down and found to contain NLRP3 by western blotting, suggesting that fluoxetine interacts with NLRP3 (Fig. 1*B*). In contrast, GAPDH was not detected in the same lysate pull-down, indicating that fluoxetine did not non-specifically bind random proteins in this assay (Fig. 1*B*). Next, a competition assay was performed by incubating the protein lysate and biotinylated fluoxetine with increasing amounts of excess unlabeled fluoxetine. Increasing ratios of unlabeled to labeled fluoxetine resulted in reduced abundance of NLRP3 detected in the precipitate (Fig. 1*B*). Ratios of 10-fold and higher excesses of unlabeled fluoxetine resulted in undetectable levels of NLRP3. These data provide support for the specificity of the presumed interaction between fluoxetine and NLRP3. Next, using an in vitro assay, we observed a direct interaction between biotinylated fluoxetine and purified recombinant myc-tagged NLRP3 protein (Fig. 1*C* and *SI Appendix*, Fig. S2). Excess amounts of unlabeled fluoxetine (5–10 fold) successfully competed against this interaction, confirming the specificity of this interaction (Fig. 1*C*). Thus, similar to CY-09 (Jiang et al., 2017), fluoxetine binds NLRP3.

### Fluoxetine binds to the NLRP3 NACHT domain in silico with predicted submicromolar affinity

Several lines of evidence point to structural mimicry of ATP, an essential ligand for NLRP3 activation, as a possible basis for the interaction of both CY-09 and fluoxetine with NLRP3. They both contain the trifluoromethyl (CF_3_) group, which, because of the strong electronegativity of fluorine, is an effective mimic for the terminal phosphate group of ATP. In addition, we have previously demonstrated that NLRP3 is inhibited by certain modified nucleoside analogs (Fowler et al., 2014), which as such also have the potential to be inhibitors of ATP binding. Thus, we assessed the hypothesis that fluoxetine could be binding in the same structural region as ADP or ATP, a site that is deeply buried in the protein at the interface of several subdomains (NBD, WHD and HD1) of the NACHT domain of NLRP3, as depicted in Fig. 2a of Sharif et al. (2019).

**Figure 2.**
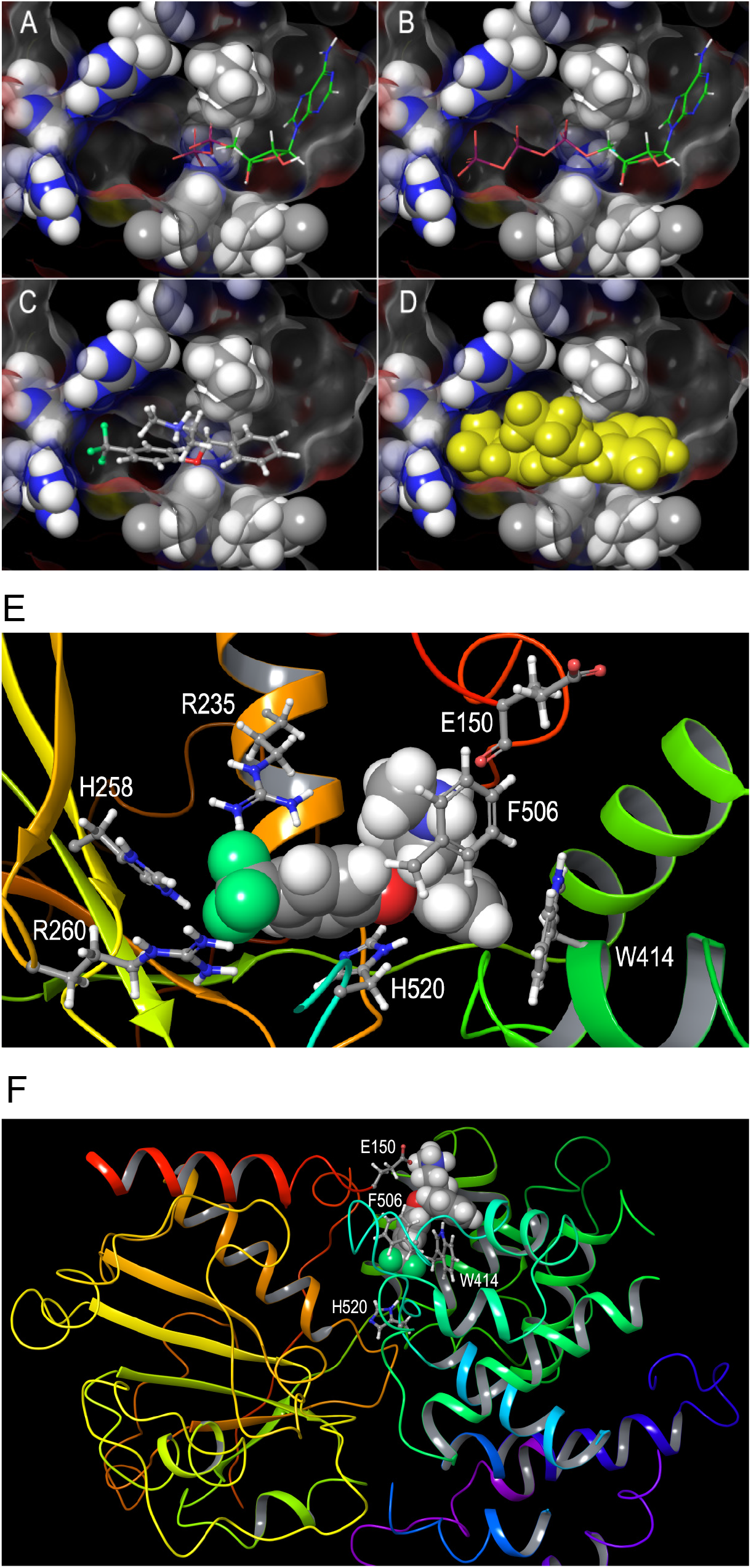
Fluoxetine binds NLRP3 in silico. (A–D) Binding modes of ADP and ATP to NLRP3 as a basis for fluoxetine binding. **A.** The fully internal cavity in which ADP is bound to the NACHT domain of NLRP3 is shown in a Z-clipped cutaway, with the internal protein surface and several key residues depicted, including several hydrophobic residues that lead to a narrowing of the cavity around the nucleotide 5’-carbon, and a cluster of basic residues (Arg235, His258 and Arg260) at the left. In the 6NPY PDB structure shown, there is a substantial void in the cavity near those basic residues. **B.** A simple rotation of the γ dihedral angle (C4’-C5’ bond) of the ribose sugar of the bound ADP, to the-*sc* orientation, along with the addition of a terminal phosphate group to make ATP, shows that a bound ATP in an otherwise identical binding mode as ADP could engage in electrostatic interactions between the terminal phosphate oxygens and the triad of basic residues mentioned above, partially filling the void in the cavity. This could represent an alternate mode of ATP binding prior to hydrolysis, possibly associated with an inactive state. **C.** The CF_3_ group of fluoxetine, with negative charges on fluorine, could act as a phosphate isostere. If it is overlapped with the terminal phosphate group as modeled in panel B, a low energy extended conformation of fluoxetine can be fitted onto the molecule of ATP shown in panel B, with the second aromatic ring of fluoxetine occupying the space of the ribose moiety. In this preliminary manual docking (see Methods), the N-methyamine sidechain of S-fluoxetine projects towards the viewer, i.e., the part of the cavity closest to the protein surface. **D.** This initial manual docking of S-fluoxetine into the nucleotide binding cavity of NLRP3 is shown with the ligand in yellow space fill rendition, showing that S-fluoxetine has the potential to fit well within the experimentally observed cavity. This hypothesis was then assessed using in silico docking methods. **E.** Best docked pose of S-fluoxetine fully inside the nucleotide binding cavity as identified by HADDOCK. The conformation of the complex shown is after an OPLS3 force field minimization. The viewpoint is slightly rotated around the Y axis relative to Fig. 2, for clarity in showing the interacting sidechains. These include the triad of basic residues mentioned in Fig. 2 (R235, H258, R260, using numbering system of 6NPY.PDB), two nitrogenous aromatic amino acids (H520 and W414), and two residues that engage the S-fluoxetine amino group in H-bonding (E150) or Pi-cation bonding (F506). The binding energy of this complex was calculated as ΔG = –8.9 Kcal/mol, using the PRODIGY-LIGAND web server (Kurkcuoglu et al., 2018; Vangone et al., 2019). A complete schematic interaction diagram for this complex is shown as *SI Appendix*, Fig. S3*A*. **F.** Alternate docked pose of S-fluoxetine only partially inside the nucleotide binding cavity. HADDOCK yielded a marginally slightly higher ranking to a cluster best represented by this complex, in which S-fluoxetine interacts with 4 of the same residues as the “fully inside” complex shown in Fig. 2*E*. These residues include E150, except the fluoxetine amino group forms a salt bridge to the glutamate side chain, rather than H-bonding to the backbone carbonyl. Trp414, Phe505 and His520 all interact with S-fluoxetine, but in different ways. This conformation could represent an intermediate state of entry of fluoxetine into the buried cavity. The third least highly ranked cluster was similar to this, except the drug molecule is inverted with CF_3_ group of fluoxetine protruding slightly from the protein. It had less favorable electrostatic binding energy than the pose shown here with the amino group interacting with the glutamate side chain.

We identified an unoccupied void in the ADP binding cavity of NLRP3 in the 6NPY cryo-EM structure (Sharif et al., 2019), that is lined on one side by basic (positively charged) residues and can accommodate the entire triphosphate of ATP when it is projected into that void via a rotation around the nucleotide C4’-C5’ bond (Fig. 2 *A* and *B*). A modeled low energy extended conformation of fluoxetine was docked in this cavity by superimposing the CF_3_ of fluoxetine on the terminal phosphate group of ATP in this pose, with the second aromatic ring of the fluoxetine molecule overlapping the ATP ribose. This orients the N-methylamine sidechain of fluoxetine up towards the “exit” of the cavity, with a good preliminary steric fit of FLX into the space (Fig. 2 *C* and *D*). Via a protein-ligand docking study using the HADDOCK web server (van Zundert et al., 2016), this mode of binding was confirmed as one of 3 clusters of docked poses in which fluoxetine was either wholly or partially inside the nucleotide binding cavity. The pose shown in Fig. 2*E*, essentially identical to the manually docked pose of Fig. 2 *C*, was the best ranked pose in the only cluster in which fluoxetine was fully inside the cavity. A schematic diagram of all of the interacting amino acid residues for this pose is shown in *SI Appendix*, Fig. S3*A*. In the two other clusters for docked fluoxetine, the fluoxetine molecule is essentially blocking the entrance to the nucleotide binding cavity. The highest ranked is shown in Fig. 2*F*, with the fluoxetine amino group projecting out from the surface of the protein. The third cluster was similar, but with the fluoxetine molecule inverted so that the CF_3_ is projecting on the outside surface of the protein. However, these “partially inside” dockings may not be accessible in the actual NLRP3 molecule, because there is an external loop of ~10 residues that is missing (disordered) in the 6NPY structure that is likely to partially occupy the region where fluoxetine is docked in these 2 clusters. So the docking shown in Fig. 2*E* is overall the most relevant. The PRODIGY LIGAND program (Kurkcuoglu et al., 2018; Vangone et al., 2019) was used to calculate the free energy of binding of fluoxetine to NLRP3, giving ΔG = −8.9 Kcal/mol, which corresponds to an inhibition constant (K_i_) of around 0.5 μM at 37°C.

### Fluoxetine inhibits NLRP3 assembly and activation

Since fluoxetine binds NLRP3, we investigated whether it disrupts inflammasome assembly, which we monitored by assessing ASC speck formation in mouse bone marrow derived macrophages (BMDMs). ASC speck formation was robustly induced by *Alu* RNA transfection but decreased by fluoxetine treatment to baseline levels (Fig. 3*A*). Next, we assessed inflammasome activation in BMDMs and immortalized human RPE cells (ARPE-19) by monitoring caspase-1 activation via western blotting. Caspase-1 cleavage was robustly induced by *Alu* RNA or B2 RNA transfection and reduced by fluoxetine treatment (Fig. 3*B*).

**Figure 3.**
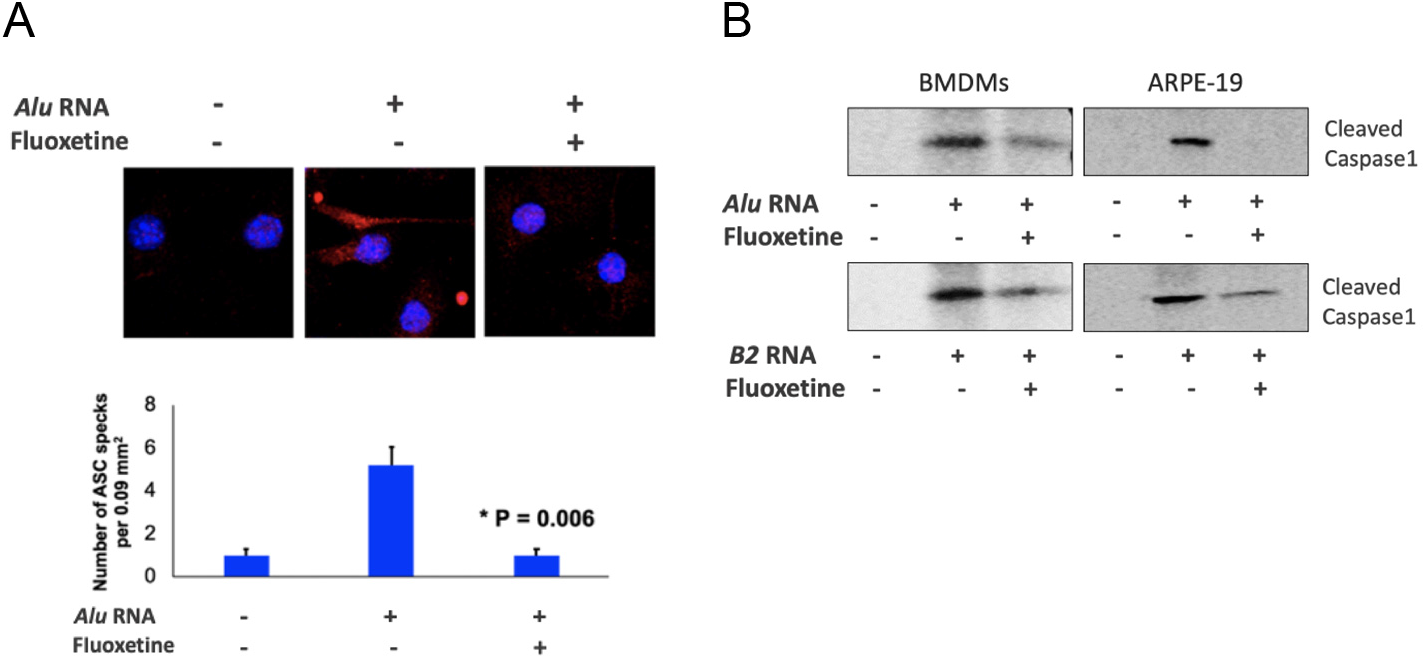
Fluoxetine inhibits SINE RNA-induced NLRP3 inflammasome assembly and activation. **(A)** Top three panels show representative immunofluorescent images of ASC specks (red circles) in wild-type mouse bone marrow-derived macrophages (BMDMs). Cell nuclei stained blue by DAPI. Bottom panel shows quantification of top panel. Mean and SEM. N = 5 per group. *P = 0.006 (*Alu* RNA + FLX vs. *Alu* RNA), Student’s two-tailed t test. FLX, fluoxetine. **(B)** Representative western blot images show *Alu* RNA induces cleavage of pro-caspase-1 (50 kDa) into active caspase-1 (20 kDa) in BMDMs (left) and ARPE-19 cells (right), and that fluoxetine (FLX) reduces caspase-1 activation in both cell types.

### Fluoxetine inhibits RPE degeneration

Next, the effects of fluoxetine and several other FDA-approved drugs belonging to various classes of antidepressants were tested in an *in vivo* mouse model of *Alu* RNA-induced RPE degeneration (Kaneko et al., 2011). RPE degeneration was assessed in two ways (Kerur et al., 2018): (1) masked grading by two readers and (2) quantitative morphometric assessment of polymegethism – the variation in the size of RPE cells. *Alu* RNA induced RPE degeneration that resembled the morphology of RPE cells in human dry AMD, whereas fluoxetine treatment conferred a profound protection against this toxicity (Fig. 4*A*). In contrast, none of the eight other anti-depressants we tested were able to inhibit RPE degeneration (Fig. 4 *A* and *B*). None of the nine anti-depressants tested induced RPE degeneration without *Alu* RNA treatment (*SI Appendix*, Fig. S4). These data demonstrate an inhibitory effect of fluoxetine in a human disease-relevant animal model and that fluoxetine exhibits a specific protective effect not possessed by multiple other FDA-approved anti-depressants.

**Figure 4.**
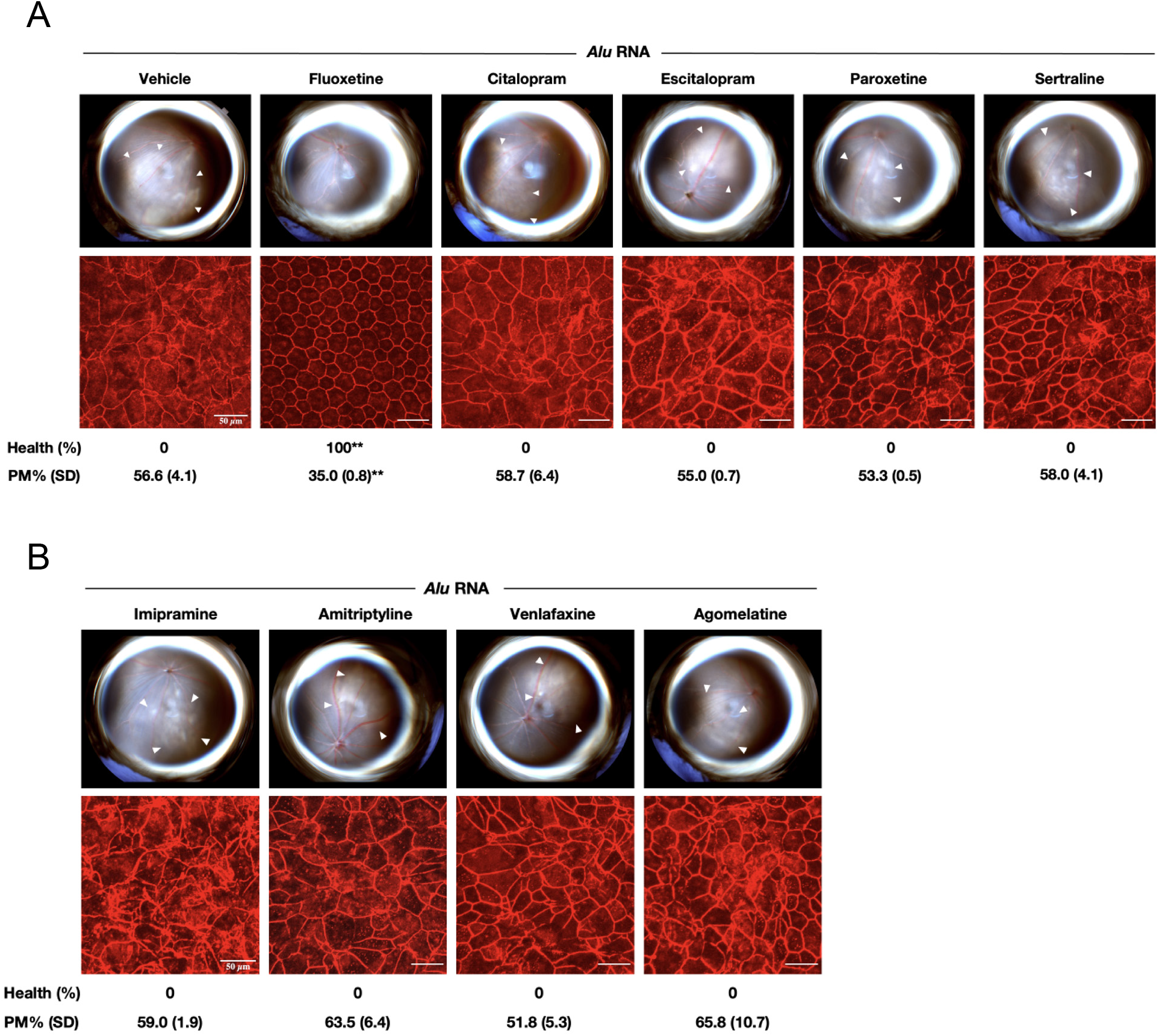
Fluoxetine inhibits *Alu* RNA-induced RPE degeneration *in vivo*. *Alu* RNA-induced RPE degeneration is blocked by fluoxetine but not by other anti-depressant drugs. (**A, B**) Subretinal administration of *Alu* RNA and intravitreous administration of selective serotonin reuptake inhibitor (SSRI) (**A**) and non-SSRI (**B**) anti-depressant drugs in wild-type mice. Fundus photographs, top row; Flat mounts stained for zonula occludens-1 (ZO-1; red), bottom row. Degeneration outlined by white arrowheads. Binary (Healthy %) and morphometric (PM, polymegethism (mean (SEM)) quantification of RPE degeneration is shown (Fisher’s exact test for binary; two-tailed t test for morphometry; ** P < 0.001). Loss of regular hexagonal cellular boundaries in ZO-1 stained flat mounts is indicative of degenerated RPE. Scale bars (50 μm).

### Fluoxetine associated with reduced development of dry AMD

The use of fluoxetine for the treatment of clinical depression for over 30 years afforded us the opportunity to assess the risk of development of dry AMD by performing a retrospective, longitudinal cohort analysis among patients aged 50 or older (the population at risk for dry AMD development). We studied the Truven Marketscan Commercial Claims database, which contains data on 90 million Americans from 2010 to 2018, and the PearlDiver Mariner database, which contains data on 15 million Americans from 2010 to 2018 (*SI Appendix*, Tables S1 and S2).

We performed Kaplan-Meier survival analyses to estimate the probability of developing dry AMD: fluoxetine use was associated with a significantly slower rate of developing dry AMD in both the Truven and PearlDiver databases (Fig. 5 *A* and *B*). Next, we performed Cox proportional hazards regression analyses to estimate the hazard of dry AMD in relation to fluoxetine use. There were similar protective associations between fluoxetine exposure and incident dry AMD in the Truven (unadjusted hazard ratio, 0.705; 95% CI, 0.675 to 0.737; P < 0.001) and the PearlDiver databases (unadjusted hazard ratio, 0.704; 95% CI, 0.639 to 0.776; P < 0.001).

**Figure 5.**
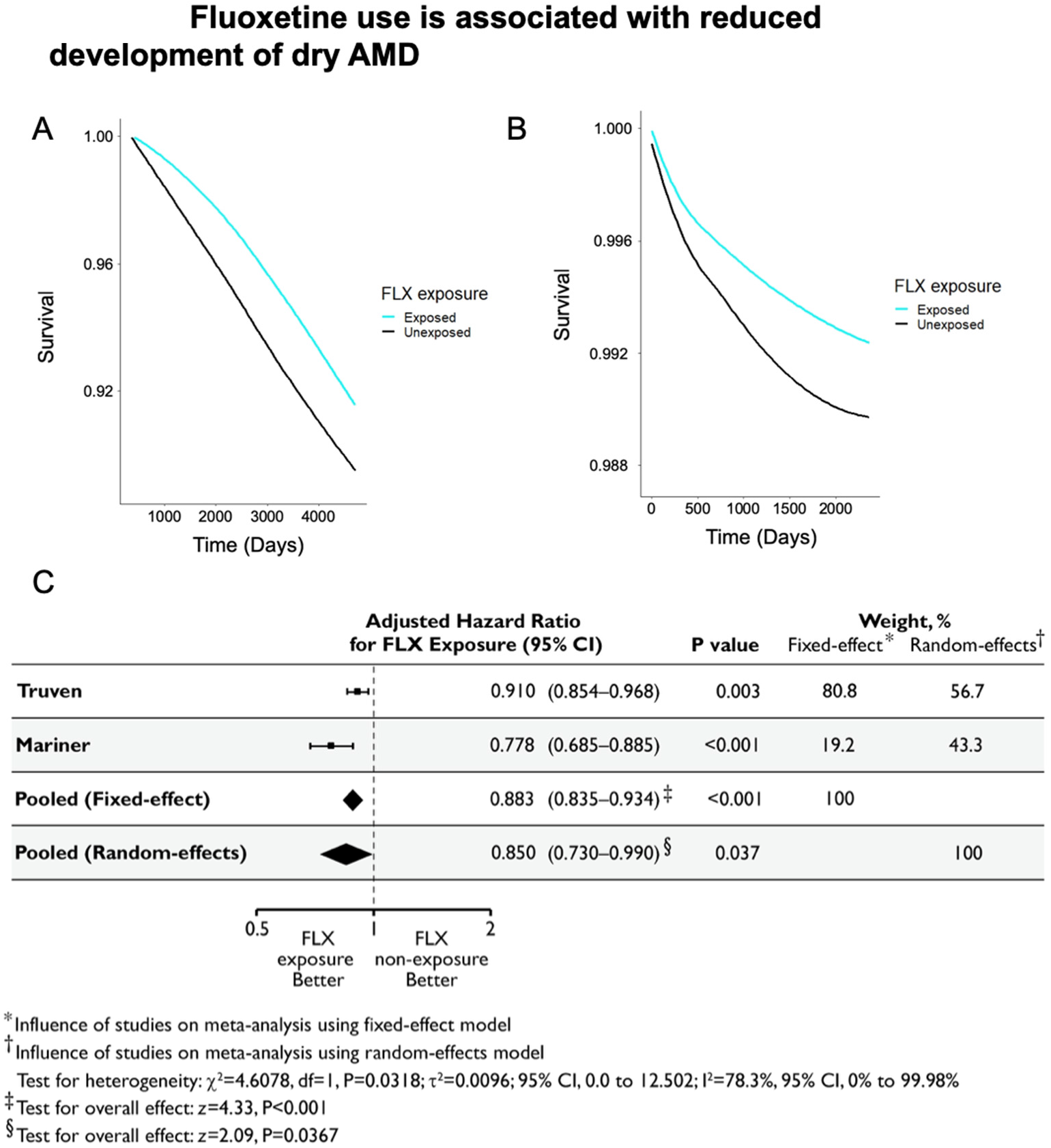
Risk of developing dry age-related macular degeneration is reduced with fluoxetine exposure. (**A, B**) Kaplan-Meier survival curves showing the probability of not developing dry age-related macular degeneration (survival) over time for subjects in the Truven Marketscan (**A**) and PearlDiver Mariner (**B**) databases (baseline characteristics in *SI Appendix*, Tables S1 and S2) based on fluoxetine (FLX) exposure or non-exposure. Difference between fluoxetine (FLX) exposure or non-exposure groups was significant (P < 0.0001 by log rank test). **(C)** Hazard ratios for developing dry age-related macular degeneration derived from propensity score-matched models (baseline characteristics in *SI Appendix*, Tables S3 and S4) adjusted for the confounding variables Methods and Materials were estimated separately for the Truven Marketscan and PearlDiver Mariner databases. Adjusted hazard ratios along with their 95% confidence intervals are shown as black lines. Diamonds show the pooled estimate of the adjusted hazard ratio and the 95% confidence intervals for meta-analyses using inverse-variance-weighted random-effects and fixed-effect models. The broken vertical line represents an adjusted hazard ratio of 1, which denotes equal risk between fluoxetine exposure and non-exposure. Horizontal bars denote 95% confidence intervals (CI). P values derived from z statistics for individual databases are reported. The estimates of heterogeneity (χ^2^), results of the statistical test of heterogeneity using the chisquare (χ^2^) test statistic and its degrees of freedom (df), and posterior probabilities of a non-beneficial effect for each model are shown below the plot. The Higgins I^2^ statistic and its 95% CI are presented. The results of the statistical tests of overall effect, the z test statistics, and corresponding P values are presented.

Patients in these databases were not randomly assigned to fluoxetine treatment; therefore, we performed propensity score matching, a causal inference approach used in observational studies (Rosenbaum & Rubin, 1983; Imai & van Dyk, 2004; Haukoos & Lewis, 2015; Ohlsson & Kendler, 2020), to assemble cohorts with similar baseline characteristics, thereby reducing possible bias in estimating treatment effects (*SI Appendix*, Tables S3 and S4). Additionally, to control for any residual covariate imbalance, we adjusted for confounders known to be associated with dry AMD: age, gender, smoking, and body mass index, as well as Charlson comorbidity index, a measure of overall health. These adjusted Cox proportional hazards regression models in the propensity-score-matched populations also revealed a protective association of fluoxetine use. In the Truven database, fluoxetine exposure was associated with a 9% reduced hazard of developing dry AMD (adjusted hazard ratio, 0.910; 95% CI, 0.854 to 0.968; P = 0.003). In the Mariner database, fluoxetine exposure was associated with a 22% reduced hazard of developing dry AMD (adjusted hazard ratio, 0.778; 95% CI, 0.685 to 0.885; P < 0.001).

Next, we estimated the combined hazard in the two databases based on an inverse-variance-weighted meta-analysis using a random-effects model. We chose this model for two reasons: (1) a substantial amount of the variance between the studies could be attributed to heterogeneity (I^2^ = 78.3%; 95% CI, 0.0% to 99.98%; P = 0.03); (2) the Truven and PearlDiver databases represent populations whose underlying true effects that are likely different (DerSimonian & Laird, 1986; Anello & Fleiss, 1995; Lau et al. 1998). The random-effects meta-analysis identified a protective effect of fluoxetine against incident dry AMD (pooled adjusted hazard ratio = 0.850; 95% CI, 0.730, 0.990; P=0.037). For completeness, we also performed a meta-analysis using a fixed-effect model; this too revealed a similar fluoxetine protective effect (Fig. 5*C*).

## DISCUSSION

Our studies demonstrate that fluoxetine directly binds NLRP3 and inhibits both NLRP3-ASC inflammasome assembly and activation. The docking results for fluoxetine are consistent with the experimental results and also show how the biotinylated analogue is able to bind, as in the two best docked poses presented here (Fig. 2 *E* and *F*), the amino group to which the linker is attached is at or nearest to the opening of the binding cavity, implying minimal disruption to the predicted mode of binding. The K_i_ of ~0.5 μM predicted by the docking simulations is compatible with the significant inhibition observed experimentally at a concentration of 10 μM. We also demonstrate that this FDA-approved drug blocks RPE degeneration in an animal model and that its use is associated with a reduced risk of developing dry AMD in humans. Collectively, these findings provide a strong rationale for launching a prospective randomized clinical trial of fluoxetine for dry AMD.

Despite numerous advances into the mechanisms of dry AMD, there is still no approved therapy for this disease. Traditional approaches to drug development can be expensive and time-consuming: on average, a new FDA-approved drug takes 10–12 years and costs $2.8 billion (present-day dollars) to develop (DiMasi et al., 2016). Our identification of the unrecognized therapeutic activity of an existing FDA-approved drug using Big Data mining, coupled with demonstrating its efficacy in a disease-relevant model could greatly accelerate and reduce the cost of drug development.

A strength of our health insurance database analyses is that findings were replicated in two independent cohorts that comprise a substantial fraction of American adults with health insurance. In addition, we adjusted for confounders and performed propensity score matching, which increases the internal validity of our conclusion. However, because there was no randomization in our study, residual confounding or selection bias might still exist. In addition, our study, like all observational health insurance claims studies, have inherent limitations in assessing the accuracy of coding and clinical phenotyping. Our studies do, however, provide a rationale for performing randomized controlled trials of fluoxetine for dry AMD, which can provide insights into causality. Demonstrating fluoxetine’s benefit for dry AMD in a prospective trial could benefit millions of patients suffering from the risk of irrecoverable blindness. It would also be interesting to determine whether fluoxetine is beneficial in other inflammasome-driven diseases such as Alzheimer’s disease, Parkinson’s disease, and diabetes (Heneka et al., 2018; Masters et al., 2011).

## Author contributions

M.A., I.A., S.N., S.W., J.M., H.L., F.P., A.V., K.B., K.M., M.S., C.I.S., B.C.W., S.R.S., E.W.T., S.S.S., and B.D.G. performed experiments or analyzed data. M.A. and B.D.G. conceived and directed the project, and wrote the paper with assistance from S.N., S.W., and I.A. All authors had the opportunity to discuss the results and comment on the manuscript.

## ACKNOWLEDGMENTS

We thank D. Robertson, G. Pattison, J. Hu, and K.A. Fox for their technical assistance. B.D.G. received support from NIH grants (R01EY028027 and R01EY031039), BrightFocus Foundation, and the Owens Family Foundation. C.I.S. received support from the NIH (R35GM119751) and the University of Virginia. The FP7 WeNMR (project# 261572), H2020 West-Life (project# 675858) and the EOSC-hub (project# 777536) European e-Infrastructure projects are acknowledged for the use of their web portals, which make use of the EGI infrastructure with the dedicated support of CESNET-MetaCloud, INFN-PADOVA, NCG-INGRID-PT, TW-NCHC, SURFsara and NIKHEF, and the additional support of the national GRID Initiatives of Belgium, France, Italy, Germany, the Netherlands, Poland, Portugal, Spain, UK, Taiwan and the US Open Science Grid. The content of this article is solely the responsibility of the authors and does not necessarily represent the official views of the National Institutes of Health. The funders had no role in study design, data collection and analysis, decision to publish, or preparation of the manuscript.

## Data availability

All data needed to evaluate the conclusions in this paper are available in the main text and *SI Appendix*.

## METHODS AND MATERIALS

### Synthesis of Biotinylated Fluoxetine

The reaction was performed in oven-dried glassware under N2. All reagents and solvents were used as commercially supplied. HPLC purification was performed using a Waters 1525 Binary HPLC pump with a 2489 UV/Vis detector. Large scale purification was done with a semi prep column (YMC-Pack ODS-A 5 μm, 250×20 mm) using a gradient of 10-95% acetonitrile containing 0.1% TFA over 47 mins in water containing 0.1% TFA. Mass spectrum was recorded using electrospray ionization mass spectrometry (ESI, Advion Expression, CMS, a single-quadrupole compact MS). Mass data is reported in units of m/z for [M+H]^+^ or [M+Na]^+^.

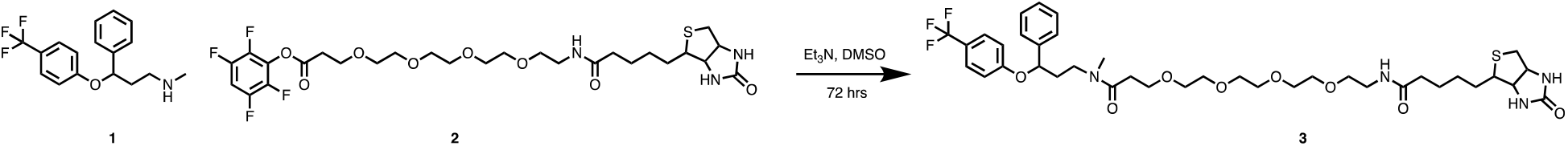

Fluoxetine (0.211 mmol, 65.5 mg, 4 eq; **1**) and TPF-PEG3-Biotin (0.052 mmol, 33.5 mg, 1 eq; **2**) were dissolved in dry DMSO (2 mL) under nitrogen atmosphere. An excess amount of triethylamine (0.1 mL, 13 eq) was added dropwise to the reaction vial. The reaction mixture was stirred vigorously under an inert atmosphere for 72 hours, followed by evaporation of the solvent and the remaining triethylamine under a high vacuum. The remaining colorless oil was dissolved in a mixture of acetonitrile and water (2 mL, 50/50 acetonitrile/water with 0.1% TFA) and purified by HPLC as described above. MS (ESI) m/z calculated for C_36_H_53_F_3_N_4_O_6_S [M+H]^+^ = 783.3, found = 783.3 (biotinylated fluoxetine; **3**).

### Purification of Myc-tagged human NLRP3 protein

To purify recombinant NLRP3 protein, HEK293T cells were transfected with plasmid encoding Myc-tagged human NLRP3 using the Lipofectamine™ 3000 transfection kit (L3000015, Thermo fisher). At 24 h after transfection, cells were collected with pre-chilled PBS and total protein was extracted using NP40 lysis buffer. The lysates were centrifuged at 12,000 rpm for 10 min at 4°C. Supernatants were incubated with anti-c-Myc magnetic beads (88842, ThermoFisher) f overnight at 4°C on rotation and then washed with lysis buffer twice. For elution of myc-tagged NLRP3 protein, incubated beads were rotated with elution buffer (50 mM Tris, pH 7.5, 500 mM NaCl, and 10 mg/ml Myc peptide (M2435, ThermoFisher)) for 60 min at room temperature. Eluted fractions were concentrated using centrifugal filters (UFC510008; Sigma). The purified proteins were validated by EZBlue™ Gel Staining (G1041, Sigma) or immunoblotting.

### Streptavidin pull-down assay

To assess the interaction of fluoxetine and NLRP3, LPS-primed THP-1 cell lysates or purified recombinant NLRP3 protein were pre-cleaned using streptavidin magnetic beads (88816, ThermoFisher) to remove nonspecific binding. These pre-treated proteins were incubated with biotinylated fluoxetine and free (unbiotinylated) fluoxetine for 1 hour on ice. Then, these samples were incubated with pre-activated streptavidin magnetic beads for overnight at 4°C with rotation. On the next day, the beads were washed with lysis buffer three times and then boiled with SDS sample buffer (LC2676, ThermoFisher) for further analysis.

### Molecular modeling and docking of fluoxetine-NLRP3 complexes

The finding that fluoxetine inhibits inflammasome activation suggests that an appropriate template for the modeling of NLRP3 should be one in an inactive state rather than an activated state. Thus we used the cryo-EM structure of Sharif et al. (2019), pdb file 6NPY, as a template for modeling NLRP3 interactions with fluoxetine. As well as a bound ADP molecule, this structure features an NLRP3 monomer bound to NEK7, which was deleted from the structure prior to any additional modifications. The hook-shaped leucine rich repeat (LRR) domain to which NEK is predominantly bound was also deleted, leaving the NACHT domain, which has the nucleotide binding domain near its center, with ADP bound in this structure (Fig. 2*A*). Fluoxetine was modeled using SybylX2.1 (Certara, Raleigh, NC), and low energy conformers identified using systematic search, with 7 rotatable bonds and electrostatics included. The distance D1 between the CF_3_ carbon and *para*-H substituent on the unsubstituted phenyl ring was monitored, enabling identification of fully extended (highest D1) vs. folded Y-shaped (lower D1) conformers. The global minimum had a low D1 and internal H-bond between an amino NH and the ether oxygen, but was too wide to fit in the nucleotide binding cavity of NLRP3. A cluster of the lowest energy fully extended conformers were less than 1.5 Kcal/mol above the global minimum, easily attainable in an induced fit interaction. In binding to a protein, the internal H-bond is also likely to be disrupted by competing intermolecular H-bonding opportunities. Thus the lowest energy extended conformation of fluoxetine was used to superimpose on the structure of ATP generated from the experimental ADP pose (as detailed in the legend to Fig. 2), with minor modifications to the methylamine side chain rotamers to minimize steric clashes in the cavity. Minimization of this all atom complex using the OPLS3 force field as implemented in Maestro version 10.7 (Schrodinger, Inc., Seattle, WA) generated the pose shown in Fig. 2 *C* and *D*. For this and subsequent docking calculations, several small gaps in the NLRP3 structure (disordered external loops) were not modeled, but capped with neutral amides, as these regions do not impinge on the core nucleotide binding cavity, which is defined by residues highlighted in red in *SI Appendix*, Fig. S3*B*.

Using that definition of the binding region of interest, docking of the S enantiomer of FLX in and around the nucleotide binding cavity of the truncated NLRP3 structure was performed using the HADDOCK2.4 web server at the University of Utrecht (van Zundert et al., 2016). HADDOCK successfully identified 3 clusters of poses for S-FLX interacting with at least some residues within the nucleotide binding cavity. The “best” conformer in one of those 3 clusters was essentially identical to the pose identified by the ligand overlap approach (Fig. 2). The PRODIGY LIGAND program from the same research group (Kurkcuoglu et al., 2018; Vangone et al., 2019) was used to calculate the free energy of binding of FLX to NLRP3 from the HADDOCK output PDB file of the docked complex.

### Cell culture studies

All cell culture experiments were compliant with University of Virginia Institutional Biosafety Committee regulations. The human retinal pigmented epithelial cell line ARPE-19 (ATCC) was maintained in Dulbecco’s Modified Eagle Medium (DMEM) supplemented with 10% fetal bovine serum (FBS) and standard antibiotics. Mouse bone marrow derived macrophages (BMDMs) were cultured in Iscove’s modified Dulbecco’s media (IMDM) with 10% FBS and 20% L929 supernatants. All cells were maintained at 37 °C in a 5% CO_2_ environment.

#### ASC speck imaging

BMDMs seeded on chambered coverslips (30,000 cells / well) for 12 h were pretreated with fluoxetine (10 μM) or 0.1% DMSO (control) for 2 h. Cells were transfected with *Alu* RNA using Lipofectamine 3000 (ThermoFisher) for 12 h. Coverslips were fixed with 2% paraformaldehyde (15 min at room temperature), washed with PBS, permeabilized, blocked with blocking buffer (PBS, 0.1% TX-100, 5% normal goat serum; 1 h at 4 °C), incubated with anti-ASC antibody (Adipogen; 1:300) with blocking buffer, and visualized with Alexa Fluor 555 (Invitrogen). Nuclei were stained with DAPI. Slides mounted using Fluoromount-G (SouthernBiotech) were imaged by confocal microscopy (Nikon A1R). The number of specks per 0.09 mm^2^ field among the mock-transfected, *Alu*-transfected, and *Alu*-transfected & fluoxetine-treated cells were compared by two-tailed Student’s t test. Mean and SEM values are presented.

#### Immunoblotting

Cells were transfected with *Alu* RNA or mock-transfected for 12 h and pre-treated with fluoxetine (10 μM) for 1 h and again after *Alu* RNA transfection. Proteins from the cell-free supernatant were precipitated by adding sodium deoxycholate (0.15% final), followed by adding TCA (7.2% final) and incubating on ice overnight. Samples were spun down at 13000g for 30 min and pellets were washed 2 times with ice-cold acetone. Precipitated proteins solubilized in 4X LDS Buffer with 2-mercaptoethanol were resolved by SDS-PAGE on Novex^®^ Tris-Glycine Gels (Invitrogen) and transferred onto Immobilon-FL PVDF membranes (Millipore). The transferred membranes were blocked with 5% nonfat dry skim milk for 1 h at room temperature and then incubated with primary antibody at 4 °C overnight. The immunoreactive bands were visualized using species-specific secondary antibodies conjugated with IRDye^®^. Blot images were captured using an Odyssey^®^ imaging system. The antibodies used were: mouse monoclonal anti-human caspase-1 antibody (AdipoGen; 1:1000) and mouse monoclonal antimouse caspase-1 antibody (AdipoGen; 1:1000).

### Animal Studies

Animal experiments were approved by the University of Virginia Institutional Animal Care and Use Committee. C57BL/6J mice (The Jackson Laboratory) were anesthetized with ketamine hydrochloride and xylazine.

#### Induction of RPE degeneration and drug treatments

*In vitro* transcribed *Alu* RNA (300 ng in 1 μl) or vehicle control (PBS) were injected subretinally. Fluoxetine, citalopram, paroxetine, sertraline, escitalopram, venlafaxine, amitriptyline, imipramine, agomelatine (all from Cayman Chemical; 1 mM in 0.5 μl) or PBS was injected into the vitreous humor 24 h before and immediately after *Alu* RNA injections. Animals were euthanized 7 days after *Alu* RNA injection, and the eyes were enucleated.

#### Assessment of RPE degeneration

RPE health was assessed by immunofluorescence staining of zonula occludens-1 (ZO-1) on RPE flat mounts. Flat mounts were fixed with 2% PFA, stained with rabbit polyclonal antibodies against mouse ZO-1 (Invitrogen; 1:100), visualized with Alexa Fluor 594 (Invitrogen), and imaged (A1R Nikon confocal microscope). Images were graded as healthy or degenerated in masked fashion. Proportions of eyes with degeneration among the various antidepressant treatment groups were compared using Fisher’s exact test. Polymegethism quantified by morphometry was compared using two-tailed t-test.

### Health Insurance Claims Database Analysis

#### Data Source

We used claims data from the Truven MarketScan Commercial Claims Database (IBM), containing healthcare claims and medication usage from the commercial insurance claims from employer-based health insurance beneficiaries from 2006 to 2018, and from the PearlDiver All Payer Claims MARINER Database (Colorado Springs), which contains healthcare claims and medication usage for persons in provider networks from 2010 to the second quarter of 2018. These de-identified data are HIPAA-compliant and were deemed by the University of Virginia Institutional Review Board (IRB) as exempt from IRB approval requirements.

#### Sample Selection

Patients were included in the analysis if they had continuous enrollment in the plan for at least 1 year and were at least 50 years of age at baseline. Individuals with pre-existing dry AMD (1 or more medical claims prior to diagnosis of depression) were excluded. Disease claims were identified by International Classification of Diseases (ICD)-9-CM and ICD-10-CM codes.

#### Independent Variable

Exposure to fluoxetine – the independent variable – was determined by whether patients filled ≥1 outpatient pharmacy prescriptions for generic or brand versions, either in sole form or as a combination medication, as identified by National Drug Codes.

#### Dependent Variable

Time to initial diagnosis of dry AMD was the dependent variable for this analysis.

#### Analyses

To analyze the risk of dry AMD between fluoxetine users and fluoxetine non-users, an adjusted Cox proportional hazards regression analysis was performed, and the hazard ratio was analyzed by the likelihood ratio test. This adjusted model included these confounding covariates known to influence dry AMD risk: age, gender, smoking, body mass index, and Charlson comorbidity index. Kaplan-Meier survival plots were analyzed by the log rank test. Statistical tests were two-sided. P values < 0.05 were considered statistically significant.

## Supplementary Information

**Figure S1.**
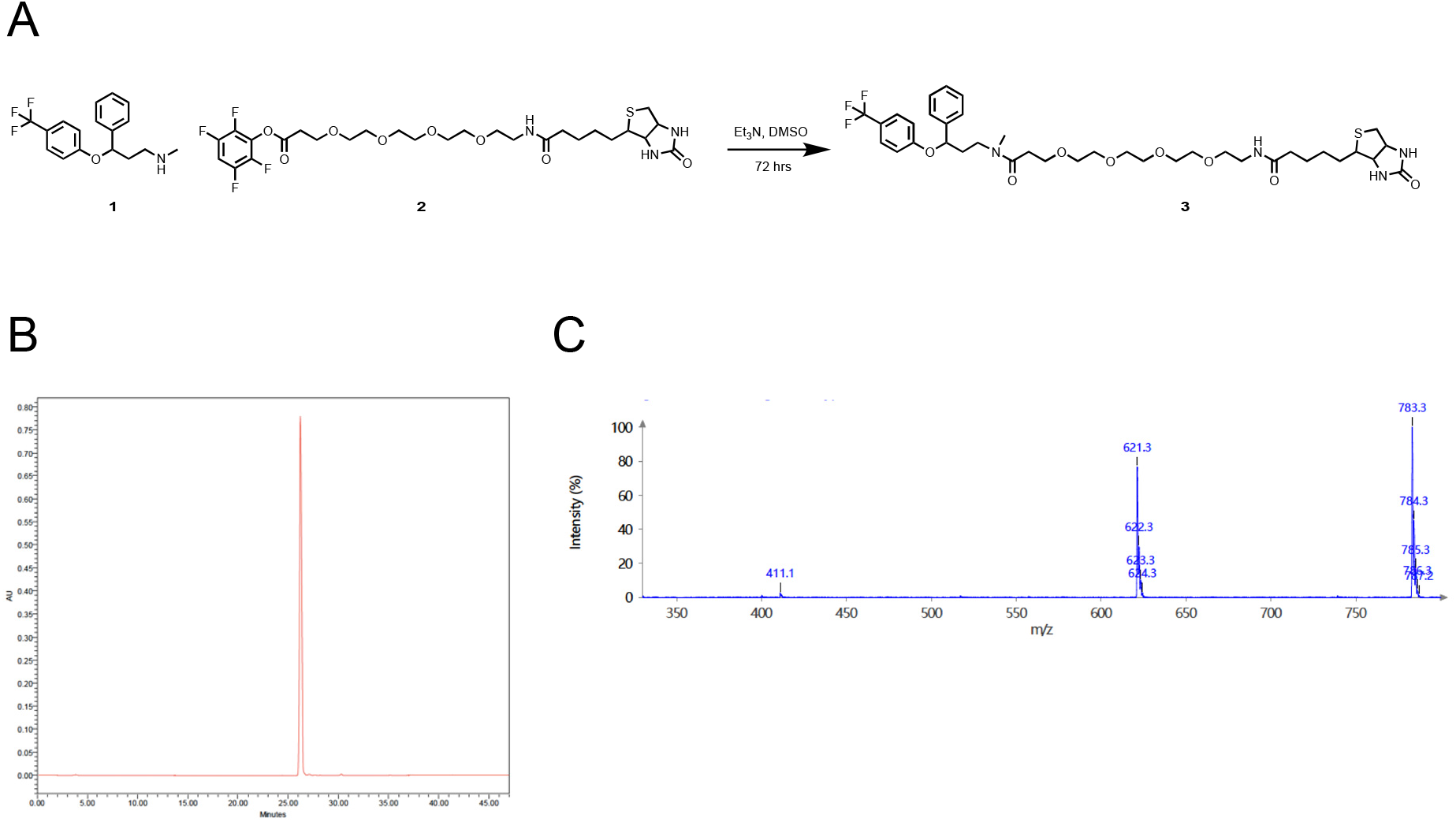
Biotin-labeled Fluoxetine. **(A)** Fluoxetine (1) and TPF-PEG3-Biotin (2), dissolved in dimethylsulfoxide (DMSO) were reacted with triethylamine (Et_3_N) to produce biotinylated fluoxetine (3). **(B)** HPLC-UV Chromatogram of biotinylated fluoxetine (3) after preparative purification. **(C)** Mass spectrum of biotinylated fluoxetine (3) detected by the ESI-MS with single quadrupole. The 621.3 m/z peak corresponds to the loss of C_7_H_4_F_3_O (161.0 m/z) fragment from biotinylated fluoxetine (3) (783.3 m/z).

**Figure S2.**
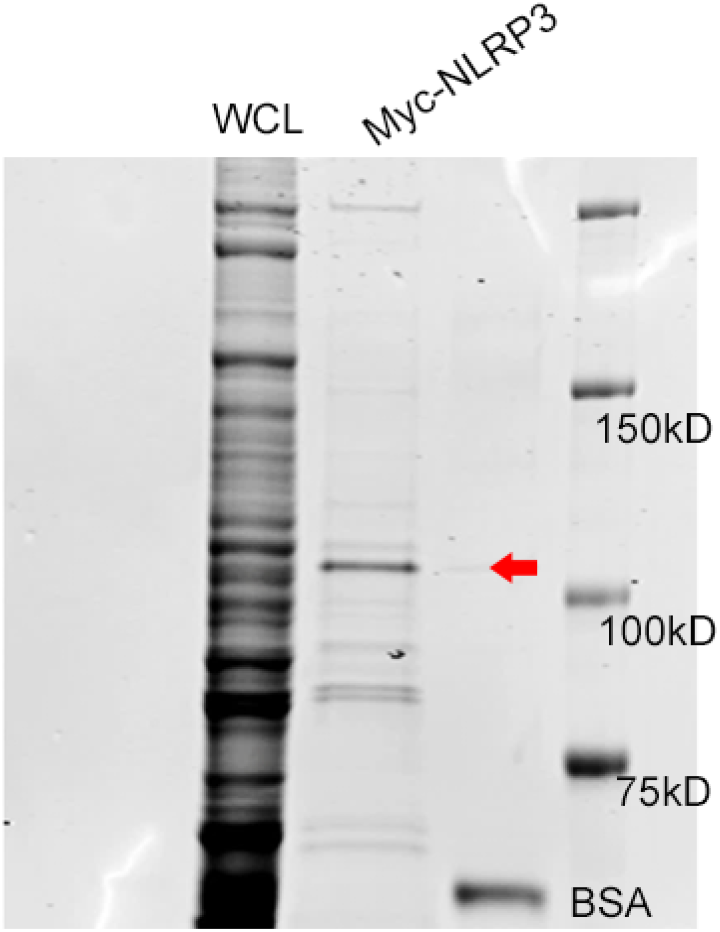
Purification of Myc-NLRP3 protein. Purification and Coomassie staining of Myc-tagged NLRP3 protein from HEK293T cells. WCL, whole cell lysates. Red arrow indicates the expected NLRP3 band.

**Fig. S3.**
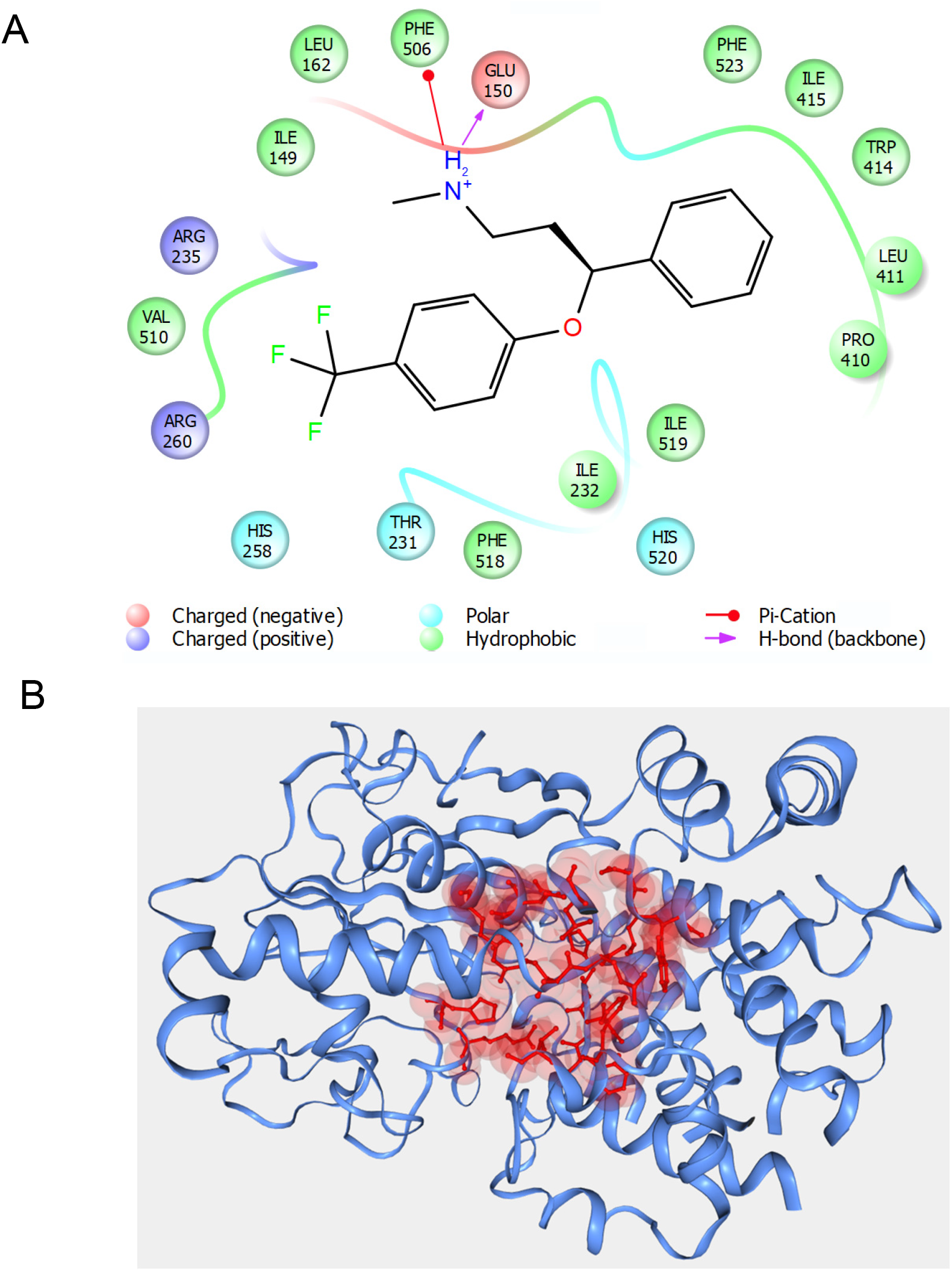
Fluoxetine NLRP3 interaction. (**A**) Complete interaction diagram for the complex of S-fluoxetine with NLRP3 shown in Fig. 2*E*. Note that there are many more interactions with hydrophobic residues that are not displayed in Fig. 2*E*. (**B**) A ribbon diagram of the entire NLRP3 NACHT domain from 6NPY showing a central regions and residues (in red) used to define the nucleotide binding cavity for the HADDOCK2.4 docking analysis.

**Fig. S4.**
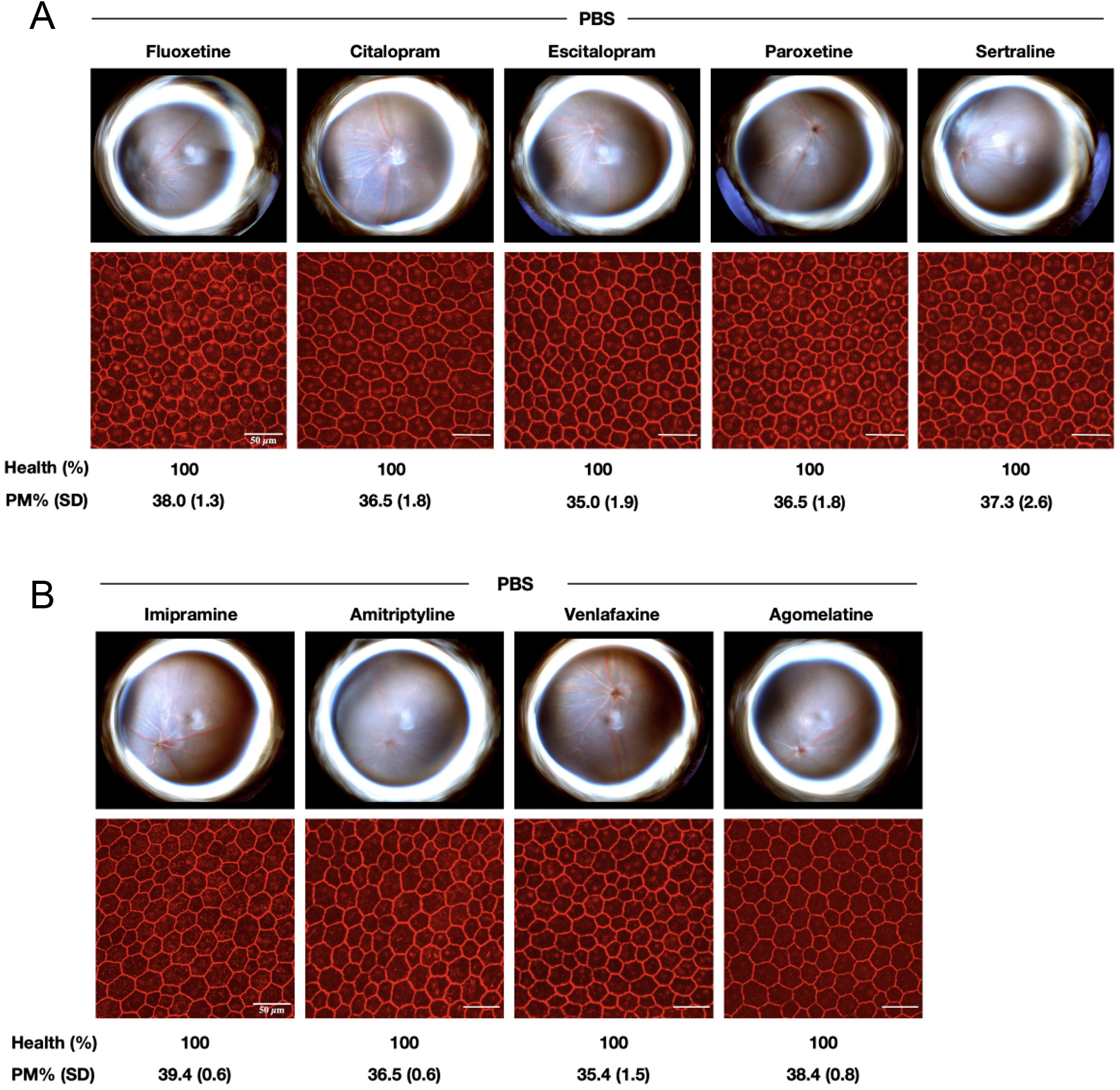
Anti-depressant drugs do not induce RPE degeneration. Anti-depressant drugs do not induce RPE degeneration. (**A, B**) Intravitreous administration of selective serotonin reuptake inhibitor (SSRI) (**A**) or non-SSRI (**B**) anti-depressant drugs in wild-type mice. Fundus photographs, top row; Flat mounts stained for zonula occludens-1 (ZO-1; red), bottom row. Degeneration outlined by white arrowheads. Binary (Healthy %) and morphometric (PM, polymegethism (mean (SEM)) quantification of RPE degeneration are shown. Scale bars (50 μm).

**Table S1.** Baseline characteristics of the study population - Truven Database. Baseline characteristics of all FLX (fluoxetine) users and a random sample of FLX non-users in the Truven Marketscan Database comprising the study population. *P-values for continuous variables are from Student t-tests and categorical from chi-square (χ^2^) tests. All statistical tests are two-sided.

| Variable |  | No record of FLX<br>N=2,215,683 | FLX Exposure<br>N=112,165 | P value* |
| --- | --- | --- | --- | --- |
| Index Age – Mean (SD) |  | 59.67(8.87) | 58.01(7.43) | <0.001 |
| Sex | Female | 66.7% | 71.6% | <0.001 |
|  | Male | 33.3% | 28.4% |  |
| Charlson comorbidity index<br>Mean (SD) |  | 0.93 (1.72) | 0.63 (1.34) | <0.001 |
| Smoking |  | 7.2% | 5.8% | <0.001 |
| Obesity |  | 8.3% | 6.2% | <0.001 |
| Duration of FLX exposure |  |  |  |  |
| Mean Days (SD) |  | 0 (0) | 607.26 (756.7) | <0.001 |

**Table S2.**
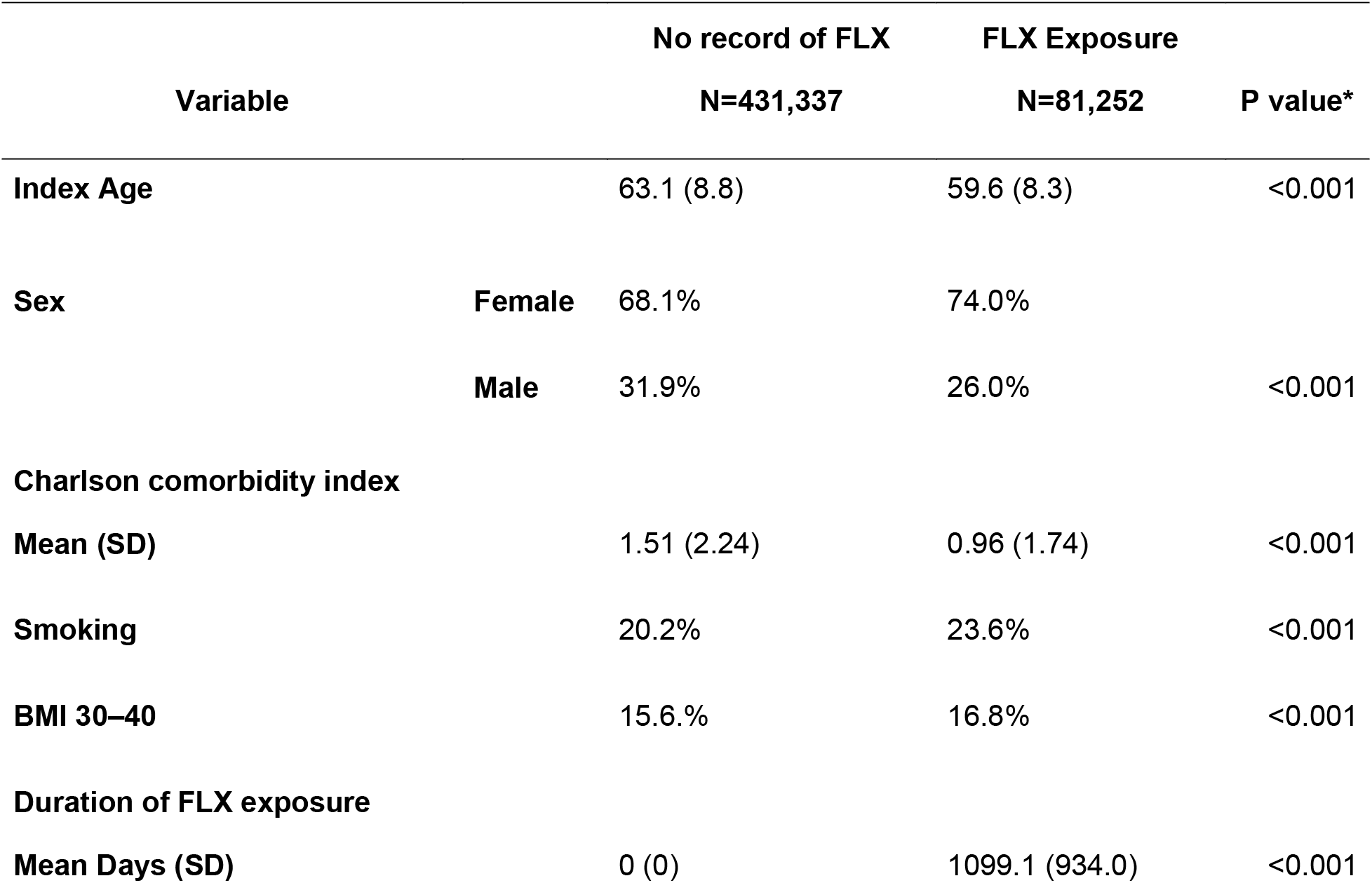
Baseline characteristics of the study population - PearlDiver Mariner Database. Baseline characteristics of all FLX (fluoxetine) users and a random sample of FLX non-users in the PearlDiver Mariner Database comprising the study population. *P-values for continuous variables are from Student t-tests and categorical from chi-square (χ^2^) tests. All statistical tests are two-sided.

**Table S3.** Baseline characteristics of propensity score matched population - Truven Database. Baseline characteristics of all FLX (fluoxetine) users and a propensity-score-matched group of FLX nonusers in the Truven Marketscan Database comprising the matched study population. *P-values for continuous variables are from Student t-tests and categorical from chi-square (χ^2^) tests. All statistical tests are two-sided.

| Variable |  | No record of FLX<br>N=112,165 | FLX Exposure<br>N=112,165 | P value* |
| --- | --- | --- | --- | --- |
| Index Age – Mean (SD) |  | 58.01(7.43) | 58.01(7.43) | 0.995 |
| Sex | Female | 71.6% | 71.6% | 0.996 |
|  | Male | 28.4.% | 28.4% |  |
| Charlson comorbidity index<br>Mean (SD) |  | 0.63(1.34) | 0.63(1.34) | 0.992 |
| Smoking |  | 5.8% | 5.8% | 0.986 |
| Obesity |  | 6.2% | 6.2% | 0.979 |
| Duration of FLX exposure |  |  |  |  |
| Mean Days (SD) |  | 0 (0) | 607.26 (756.7) | <0.001 |

**Table S4.**
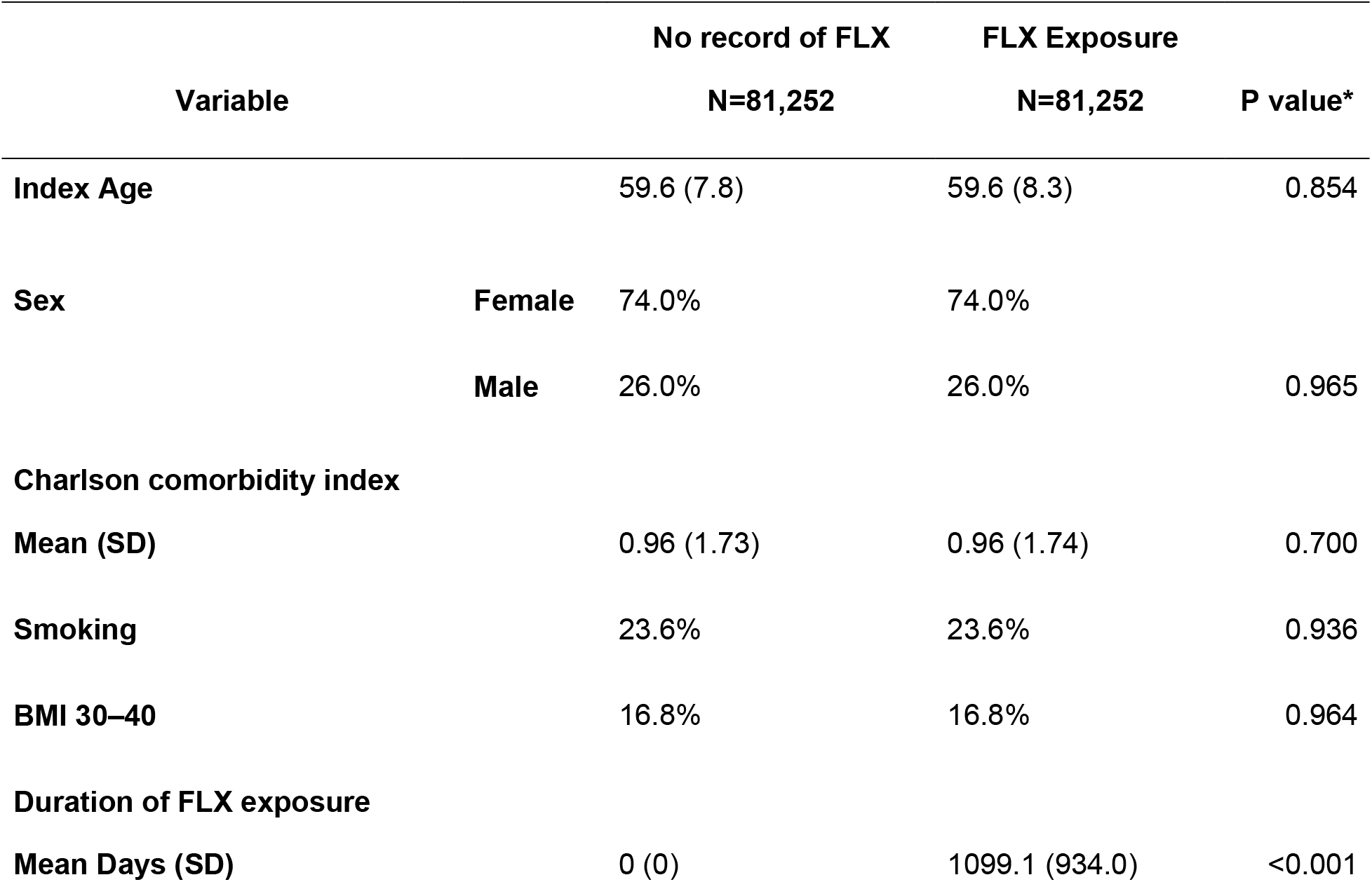
Baseline characteristics of propensity score matched population - PearlDiver Mariner Database. Baseline characteristics of all FLX (fluoxetine) users and a propensity-score-matched group of FLX nonusers in the PearlDiver Mariner Database comprising the matched study population. *P-values for continuous variables are from Student t-tests and categorical from chi-square (χ^2^) tests. All statistical tests are two-sided.

